# A dynamical model of TCR*β* gene recombination: Coupling the initiation of D*β*-J*β* rearrangement to TCR*β* allelic exclusion

**DOI:** 10.1101/200444

**Authors:** Sébastien Jaeger, Ricardo Lima, Arnaud Meyroneinc, Marie Bonnet, Edgardo Ugalde, Pierre Ferrier

**Affiliations:** Centre d’Immunologie de Marseille-Luminy, Aix Marseille Université, CNRS, INSERM, CIML, Marseille, France; Centre de Physique Théorique, Aix Marseille Université, Université de Toulon, CNRS, CPT, Marseille, France; Departamento de Matemáticas, Instituto Venezolano de Investigaciones Científicas, Caracas, Venezuela; Instituto de Física, Universidad Autónoma de San Luis Potosí, San Luis Potosí, México

## Abstract

One paradigm of random monoallelic gene expression is that of T-cell receptor (TCR)*β* allelic exclusion in T lymphocytes. However, the dynamics that sustain asymmetric choice in TCR*β* dual allele usage and the production of TCR*β* monoallelic expressing T-cells remain poorly understood. Here, we develop a computational model to explore a scheme of TCR*β* allelic exclusion based on the stochastic initiation of DNA rearrangement [V(D)J recombination] at homologous alleles in T-cell progenitors, and thus account for the genotypic profiles typically associated with allelic exclusion in differentiated T-cells. Disturbances in these dynamics at the level of an individual allele have limited consequences on these pro1les, robust feature of the system that is underscored by our simulations. Our study predicts a biological system in which locus-specific, prime epigenetic allelic activation effects set the stage to both optimize the production of TCR*β* allelically excluded T-cells and curtail the emergence of their allelically included counterparts.

## Introduction

In developing T and B lymphocytes, T-cell receptor (TCR) and immunoglobulin (Ig) genes are assembled from separate variable (V), diversity (D), and joining (J) gene segments in a process called V(D)J recombination (Schatz and Swanson, 2011; Alt et al., 2013). This biochemical reaction relies on the lymphoid-specific RAG1 and RAG2 proteins. The RAG1/2 machinery cleaves DNA at asymmetric recombination signal sequences (12-and 23-RSS) that flank each pair of rearrangeable gene segments. The resultant DNA double-stranded breaks (DSBs) are repaired by components of the non-homologous end joining (NHEJ) pathway to yield coding-and signal-joints (CJs and SJs). Coding joints often display limited deletion and/or addition of nucleotides. Randomization of CJ sequences, while contributing to immune cell diversity, ultimately only yields, on average, one third of productively (in-frame) assembled TCR/Ig genes.

V(D)J recombination obeys genetic programs that integrate signaling cues from generated receptors to sustain cell development in distinct T-and B-cell lineages. For example, during *αβ*T-cell development in the mouse thymus, TCR*β* gene recombination occurs in CD4/CD8 double negative (DN)2 and DN3 evolutionary thymocytes, with D*β*-to-J*β* rearrangement preceding V*β*-to-DJ*β* joining. Expression of a productively rearranged V*β*DJ*β* CJ (hereafter VDJ+) and formation of a primary receptor, namely the pre-TCR, triggers further differentiation into DN4 cells and subsequently CD4/CD8 double positive (DP) cells. This developmental shift is known as *β*-selection. V(D)J recombination is arrested during this period of cell differentiation. It resumes in DP cells by selectively targeting the TCR*α* locus to achieve V*α*-to-J*α* joining, followed by further selection events involving the completed *αβ*TCR. Work done by many groups over the past three decades has revealed multiple molecular levels involved in regulating these dynamics, including RAG1/2 expression, chromatin remodeling, nuclear positioning and 3D-folding of TCR/Ig loci, and occasionally, restricted 12/23-RSS pairing usage (Schatz and Swanson, 2011; Alt et al., 2013).

Individual T/B lymphocytes operate under clonal selection, and usually display TCR/Ig receptor components from only one of two chromosomal alleles, a property termed allelic exclusion (AE). For TCR*β* (as well as *Igh* and *Iglκ*/*λ*) gene products, AE primarily results from contrasting rearrangements at corresponding alleles (Khor and Sleckman, 2002). Thus, the most common TCR*β* genotypes in *αβ*T-cells is a VDJ+ on one allele, with either an intermediate DJ*β* (DJ) CJ or a non-productive V*β*DJ*β* (VDJ-) CJ on the other allele, in a ~[50%-60%]/[50%-40%] ratio (so-called 60/40 ratio), respectively. The prevailing theory for the interpretation of these profiles relies upon a regulated model wherein following biallelic DJ*β* joining and the assembly of a primary VDJ+, AE prevents further V*β* rearrangement *via* inhibitory signaling from the newly formed pre-TCR (Khor and Sleckman, 2002; Brady et al., 2010a). This scenario is supported by a wealth of experimental data, including the finding that engineered deficiencies in pre-TCR formation or pre-TCR-mediated signaling in developing T-cells result in the disruption of TCR*β* AE (von Boehmer et al., 1998).

AE completion is thought to consist of two functional phases: an initiation phase of monoallelic activation to prevent V gene rearrangements from occurring simultaneously on both alleles, and a maintenance phase of feedback inhibition to suppress continued rearrangement (Gorman and Alt, 1998). Although the latter phase is sufficiently well defined in developing T-and B-cells to rally community consensus, cracking the code of monoallelic activation still represents a topical and controversial subject. Indeed, common interpretations consider no less than three types of distinct scenarios, including: (*i*) clonal allelic predetermination on a fifty-per-cent chance basis, associated with differential replication timing and suppressive pericentromeric recruitment of the late-replicating allele (Mostoslavsky et al., 2001; Levin-Klein and Bergman, 2014); (*ii*) interchromosomal transmodulation, possibly involving dual allele pairing and ensuing regulation of monoallelic DSB cleavage (Hewitt et al., 2009; Chaumeil and Skok, 2013); or (*iii*) equal biallelic inefficiency in recombination *via* stochastic transcription or association of both alleles with the nuclear lamina (Liang et al., 2004; Mostoslavsky et al., 2004; Schlimgen et al., 2008; Murre, 2008). In addition, none of these scenarios explicitly meets a number of seemingly opposing specifications that - alongside the completion of ~60/40 [VDJ+/DJ]/[VDJ-/VDJ+] AE profiles - are distinctive to this unique form of monoallelic gene expression. For instance, it remains unclear how the mechanisms suggested by the proposed scenarios can in their simplest forms proceed differently in the case of V-and D/J-containing *cis*-domains at TCR*β*/Igh tripartite V-D-J loci (allelically excluded *versus* allelically included, respectively). Likewise, in the context of scenario (*i*) or (*iii*) particularly, it is uncertain how the system manages to proceed efficiently with biallelic usage as suggested by the frequent appearance of VDJ-/VDJ+ cells [*i.e.* following the production of a primary VDJ-joint, how could the suppressed, pericentromeric recruited allele be steadily reactivated - (scenario (*i*)); or, how could the two alleles be used in a statistically significant manner assuming a necessarily low probability *p* of individual allele usage to efficiently prevent allelic inclusion - (scenario (*iii*))?]. Finally, none of these scenarios takes into account the real fact that aside from the two prominent VDJ+/DJ and VDJ-/VDJ+ genotypes, less common TCR*β* profiles, comprising either two distinct VDJ+ or a VDJ+ and a non rearranged germline (GL) allele, are also observed in low but significant fractions of *αβ*T-cells (Khor and Sleckman, 2002; Brady et al., 2010a). In short, despite years of intense investigations, the initiation phase of AE remains a puzzling issue [also see Gorman and Alt (1998); Mostoslavsky et al.(2004); Vettermann and Schlissel(2010); Outters et al.(2015)].

One strategy for the difficult task of investigating lymphoid cell development is to use modeling and simulation [*e.g.* Claverie and Langman (1984); Shlomchik et al. (1998); Piper et al. (1999); Sepulveda et al.(2005); Volpe and Kepler (2008); Vibert and Thomas-Vaslin (2017); Krueger et al. (2017)]. In this way, we have previously used Markov chain-based modeling to address the afore-mentioned issues, and thus demonstrated that TCR*β* AE could be enforced by combining three separate concepts - allele independency, probabilistic behaviors and regulated feedback - on a dynamical basis, with all the cell subsets that emerge from the TCR*β* assembly process now incorporated into a unique distribution (Farcot et al., 2010). Transition parameters implied that V*β*-to-DJ*β* joining occurs over a longer time frame in comparison to either the D*β*-to-J*β* rearranging step or the feedback closing step (Farcot et al., 2010). Though these findings seem to support the widespread notion that the initiation of TCR*β* AE is dependent solely and exclusively on V gene recombination, paradoxical results led us to consider the new and unconventional idea that it might instead primarily depend on probabilistic events that lead to the onset of D*β*-to-J*β* rearrangement and that are perhaps overlooked by our Markov-chain-based modeling. Specifically, the period of time to achieve D*β*-to-J*β* recombination turned out to be overly brief when based on allotted Markovian times (Farcot et al., 2010). However, this initial step is in fact thought to require intricate epigenetic changes during the regional conversion from a repressive to an active chromatin status (Sikes and Oltz, 2012; Jaeger et al., 2013). Based on the compelling evidence that epigenetics contributes towards the modulation of cellular plasticity in development (Pujadas and Feinberg, 2012), we wished to explore the possibility that, in establishing the chromosomal landscape required for the onset of D*β*-to-J*β* rearrangement, noisy kinetics significantly contribute towards promoting TCR*β* AE. Our mathematical simulations presented here provide strong support for this original hypothesis.

## Results

### Modeling the dynamics of TCR*β* recombination in single developing T-cells

In DN thymocytes, TCR*β* gene activation for recombination depends on cooperative outputs from *cis*-regulatory elements, including the transcriptional enhancer E*β* and D*β*-flanking promoters pD*β*s (Sikes and Oltz, 2012). At this stage, attributes of relaxed chromatin (Pol-II-generated GL transcripts, CpG demethylation, active histone marks, positioned arrays of nucleosomes) are evident in the genomic regions that encompass the D*β* and J*β* gene segments (Morshead et al., 2003; Kondilis-Mangum et al., 2010; Zacarias-Cabeza et al., 2015). These permit the local recruitment of RAG machinery to form a so-called recombination center (Ji et al., 2010b), and also may optimize lineage-specific release of TCR*β* alleles from suppressive peripheral subnuclear positioning (Schlimgen et al., 2008; Chan et al., 2013). In mammalian cells, combinatorial epigenetic modulations are a major component of developmental, allele-specific plasticity and nuclear reorganization, and a source of intrinsic noise in gene expression (Pujadas and Feinberg, 2012; Eckersley-Maslin and Spector, 2014; Therizols et al., 2014; Soshnev et al., 2016). For genes whose activation has to reach a trigger threshold level of epigenetic changes, noise accumulated during these processes may lead to substantial interallelic/cell-to-cell variability in the time required to reach the target level (Pujadas and Feinberg, 2012; Eckersley-Maslin and Spector, 2014). Along these lines, we hypothesize that initial cross-regulatory interactions at TCR*β* alleles likewise proceed *via* threshold kinetics towards establishing the proper chromosomal landscape required for the onset of D*β*-to-J*β* rearrangement, and thus, over time, contribute to probabilistic interallelic behaviors and cell-to-cell variability.

Mathematically, the problem of determining the time required for a stochastic process to reach a certain value falls into the category of first-passage time (FPT) problems (van Kampen, 2007). However, although our current assumption primarily concentrates stochasticity on the epigenetic allele-specific activation events that are mandatory to the onset of TCR*β* recombination, it is likely that all following steps in the developmental process, including D*β*-to-J*β* and V*β*-to-DJ*β* recombination and feedback inhibition, also obey probabilistic rules, albeit, we surmise, to lower magnitudes. To ensure the simulation of all of the above conditions, we devised a model system based on discrete time modeling (for comprehensive mathematics, *SI Appendix, section S1*/*Methods*), which altogether defines a rule-based framework for timely progression of V(D)J recombination at dual TCR*β* alleles in single developing T-cells. In this system, asynchrony that is introduced during the initiation phase is propagated along the whole developmental process until an asymptotic status is achieved. Importantly however, as a corollary of discretization, delay effects [<1 time unit (*t.u.*) in length and which can therefore be overlooked with respect to overall stochasticity as a first approximation] do in fact take place during every single step in the mathematical model [*e.g.* Coutinho et al. (2006)]. Thus, the selected formalism offers a double advantage: it keeps free-parameterization at a minimum for formal challenging of a dominance of prime activation events at low risk of overfitting, and yet tolerates a lower degree of freedom during each of the subsequent steps that complete the dynamics. Specifically, our system is made up of pairs of three symmetrical, interacting variables-one triplet set for each TCR*β* allele. In line with our prime hypothesis, the first pair [(*ω*_1_, *ω*_2_); equation (1)] are Boolean input variables, which compute the stochastic transition of GL TCR*β* alleles from a recombination-incompetent stage (*ω*_*i*_=0) to a status that is primed for the onset of TCR*β* gene recombination (*ω*_*i*_=1), *i.e.* up to the precise time point at which D*β*-to-J*β* recombination is initiated (each *ω*_1_ and *ω*_2_ initialized to 0 at *t*=0). At each and every time step, there is a probability *p*_*i*_ for *ω*_*i*_ to switch from 0 to 1. Thus, *p* represents the one major stochastic parameter of our model with, typically, *p*_1_=*p*_2_=*p*. The next two pairs [(*x*_1_, *x*_2_ and *y*_1_, *y*_2_); equations (2) and (3)] are state variables, which represent the current status in the progression of each individual allele’s D*β*-to-J*β* and subsequent V*β*-to-DJ*β* rearrangements, respectively.

**Figure 1.**
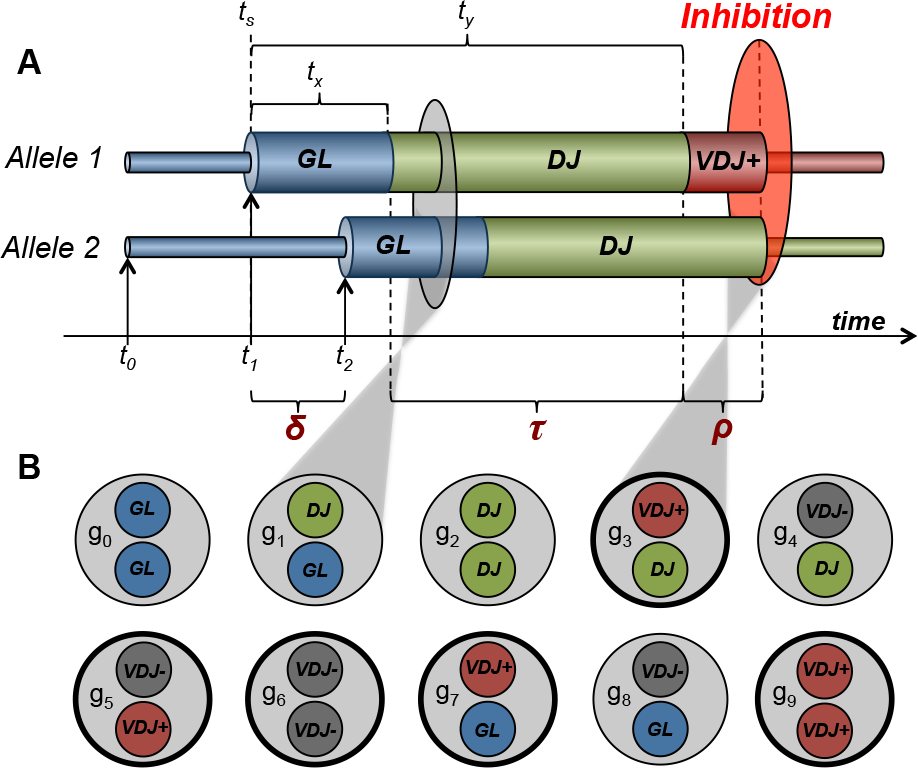
Dynamical modeling of gene rearrangement at TCR*β* alleles in single developing T-cells. (**A**) Schematic overview of one rearrangement scenario leading to the VDJ+/DJ genotype *via* probabilistic initiation of D-to-J recombination up to feedback inhibition. Thin and thick bars correspond to developmental allelic structures refractory to and compulsory for the onset of TCR*β* gene recombination, respectively. GL, unrearranged alleles; DJ, VDJ, D-to-J-and V-to-DJ-rearranged alleles, respectively. See text for the definition of the indicated time points *t* (*t*_0_, *t*_s_, *t*_1_, *t*_2_, *t*_*x*_, and *t*_*y*_) and related parameter values *δ*, *τ*, and *ρ*. (**B**) All ten possible cell arrangements in terms of TCR*β* allele configuration (denoted *g*_0_ to *g*_9_) are shown. Bold circles correspond to asymptotic cell states; examples of intermediate (*g*_1_) and terminal (*g*_3_) cell states from the rearrangement scenario depicted in (**A**) are highlighted.

An additional random variable (*є*_*i*=1,2_), incorporated into state variables *y*_*i*_, features the in-frame *versus* out-of-frame probabilistic outcomes. In the in-frame situation, the dynamics are interrupted after a fixed period of time (*ρ* ≥ 1), which corresponds to the interval that is required to achieve feedback inhibition. Conversely, in the alternative, out-of-frame, VDJ-event, progression of the rearrangement sequence continues on the opposite allele. Based on available data on TCR*β* gene rearrangement (Aifantis et al., 1997; Senoo et al., 2003; Khor and Sleckman, 2005), we considered that it was not essential to take into account either the scenario of a pre-established time window that would prematurely remove cells that had not yet used the opportunities to achieve a VDJ+ (*e.g.* V*β*DJ*β*-/DJ*β* cells) or, separately, the scenario of a significant fraction of VDJ+/VDJ+ cells arising due to the feedback mechanism not being triggered by the first rearrangement. Both scenarios would ultimately result in predicted data, including increased and decreased 60/40 ratios, re-spectively, that diverge from experimentally determined values (*SI Appendix, section S2*). Likewise, we did not consider extra variables featuring episodes of cellular division, as these mostly take place prior to D*β*-to-J*β*, and after V*β*-to-DJ*β* recombination (Ciofani et al., 2004; Masuda et al., 2007).

The dynamic rules of our model system are schematically depicted in Fig. 1A. Time *t* relates to an initial condition *t*=*t*_0_ when, in any given progenitor T-cell, all four state variables display their base values. Further time points of significance are defined as *t*_*start*_ (*t_*s*_), *δ* and *ρ*, respectively [equations (5) and (6)]. Briefly, *t*_*start*_ marks the time at which a D-to-J recombination reaction begins on the earlier-rearranging allele *i* (as one *ω*_*i*_ switches from 0 to 1 for the first time), while *δ* defines the time lag between this initial event on the two homologous alleles, specific to the given cell [geometric distribution of *δ*s directly derived from stochastic parameters *p*_*i*_, equations (10)-(13); also see *Fig. S1* for graphical representations of the theoretical distribution and partition fonction of *δ*]. For each allele, it takes *t*_*x*_ and *t*_*y*_ > *t*_*x*_* to achieve D-to-J and V-to-DJ recombination, respectively. Completion of each event corresponds to the point when *x*_*i*_ (resp. *y*_*i*_) exceeds the threshold *T*_*x*_ (resp. *T*_*y*_). Thus, the first-rearranging allele (Fig. 1A, *Allele 1*, initiating recombination time *t*_1_) completes D-to-J recombination and V-to-DJ recombination at time *t*_*s*_ + *t*_*x*_ and *t*_*s*_ + *t*_*y*_, respectively. At the remaining allele (*Allele 2*, initiating time *t*_2_), all timings are delayed by the cell-specific interval *δ*. The parameter *τ* = *t*_*y*_−*t*_*x*_ measures the whole period of time necessary for a newly generated DJ-rearranged allele to complete V-to-DJ joining. If a VDJ+ is produced, the recombination process will stop on the opposite allele at time *t*_*s*_ + *t*_*y*_ + *ρ*, where *ρ* defines the time interval required for feedback inhibition. Without loss of generality, we alternatively considered that the process also arrests if both TCR*β* alleles have achieved VDJ-rearrangements. Overall, the above parameters account for all evolutionary paths that a single cell may possibly follow when sequentially switching from one defined TCR*β* dual allele configuration to the next (out of a total of 10, denoted *g*_0_−*g*_9_; Fig. 1B).

Based on the relative values of *δ*, *τ*, and *ρ* and outcomes from *є*_*i*=1,2_, six timeline scenarios of single-cell histories are anticipated, commensurate with all orders of possible combinations (Fig. 2). Timeline scenarios cluster into three classes, each containing histories endowed with an equal probability of reaching identical dual allele configurations (*ψ* _1_, *ψ*_2_, *ψ*_3_, bordered by unique colors in Fig. 2). The value of *δ* relative to *ρ* and *ρ* + *τ* defines the class to which the individual cell belongs: *ψ*_1_ (in cases where *δ* ≤ *ρ*; pertaining to the VDJ+/VDJ-*and* VDJ+/VDJ+ cell fates); or *ψ*_2_ (*ρ* < *δ* ≤ *ρ* + *τ*; including the VDJ+/VDJ-and VDJ+/DJ cell fates); or *ψ*_3_ (*δ* > *ρ*+*τ*; comprising the VDJ+/VDJ-and VDJ+/GL cell fates). These scenarios of cell histories differentiate our dynamical model from those based on a biphasic view/instructive mode of AE initiation [which is that only following a non-productive V-to-(D)J prime allele rearrangement would the opposite allele be subsequently recombined, *e.g.* Chaumeil and Skok (2013); Levin-Klein and Bergman (2014)], which solely considers a ‘*ψ*_2_-like’ distribution of asymptotic outcomes (or ‘*ψ*_3_-like’, in case of a one-step V-J junctional event).

### Modeling the dynamics of TCR*β* genotype distributions in developing T-cells

In the model system we describe, individual cells and developmental cell fates are independent. Therefore, addressing the dynamics of TCR*β* genotypic changes in separate subsets of developing T-cells is straightforward, and is accomplished by summing up the relative fractions of single-cell profiles shown in figure 2. To begin with, we modeled the genotype evolution within a set of progenitors that concurrently engage in TCR*β* gene recombination. In this case of synchronously developing cells, the distribution of TCR*β* genotypes at any time *t* during the recombination process can be viewed as a vector 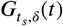 in the space of linear combinations of ten possible *g*_*i*_ subsets ( *i* = 0,…,9) [*SI Appendix, S1*/equations (8)-(14)]. Cells that leave the simulated developmental system independently of TCR*β* rearrangement outcomes (*e.g.* towards the *γδ* T-cell lineage) should have little effect on these distributions, in so far as relative proportions are considered, as they are anticipated to be evenly split among the various subsets at each time step.

**Figure 2.**
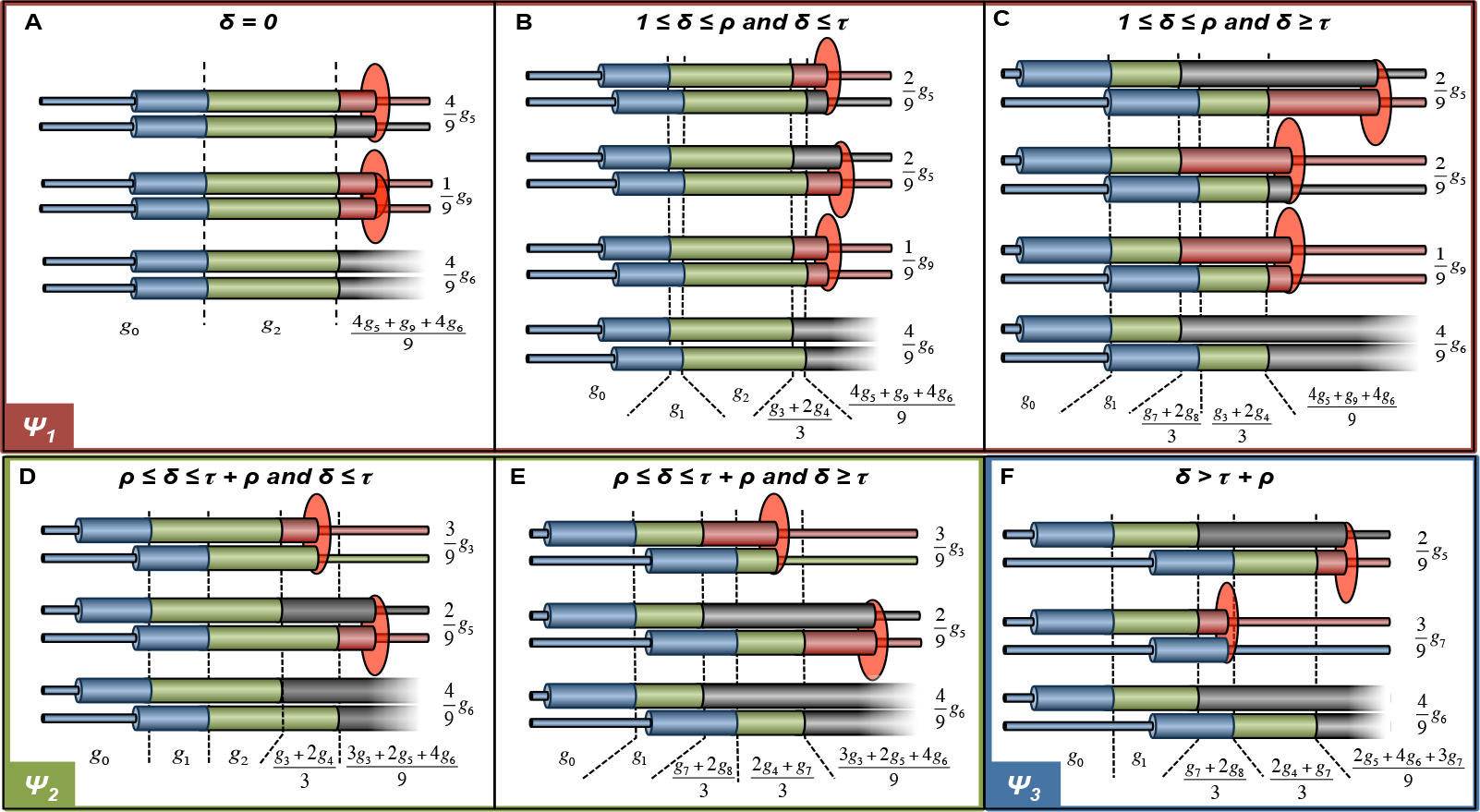
Timeline scenarios of TCR*β* allelic rearrangements. Relevant histories of single T-cells or, alternatively (see text), of a synchronously developing T-cell subpopulation are shown, depending on relative *δ*, *ρ* and *τ* values and the one-third ‘*in frame*’ to two-thirds ‘*out of frame*’ rule of V-to-DJ rearrangement outcomes. The annotated fractions indicate the expected, time-scaled probabilities (or distributions for a cell population) of the corresponding *g*_0_ to *g*_9_ genomic con1gurations, with the respective probabilities (or distributions) of each final state at asymptotic regimen shown on the right. Lighter lines correspond to cells ending TCR*β* gene recombination in the VDJ-/VDJ-(*g*_6_) con1guration, doomed to disappear from the developmental process. Cell sets sharing the identical final distributions are clustered into the *ψ*_1_, *ψ*_2_, and *ψ*_3_ subclasses (gathered by red, green and blue borders, respectively), depending on explicit rules as indicated at the top of the figure. When using model calibration such as defined in the *Results*, the respective distribution between the three sub-classes is: 9% *ψ*_1_, 87.9% *ψ*_2_ and 3.1% *ψ*_3_. Note that cell behaviors, as depicted in panel (**A**) which displays developing single cells with *δ* = 0, are not necessarily negligible in terms of cell representation since both the few cells displaying no ‘time gap’ between the two rearranging alleles and those with a *δ* < 1 *t.u.* are included.

With the notable exception of early-stage thymic embryogenesis, the DN cell compartment does not typically accommodate a unique cohort of synchronously differentiating T-cells (Ramond et al., 2014). Rather, it is supplied with TCR*β*-unrearranged cell precursors, and is drained of advanced cells, including TCR*β*-rearranged (productively or not) thymocytes. Our mathematical model can be applied to these cellular inflow/outflow situations by considering cell-subsets as before, while also having distinct *t*_0_. To integrate cell departure, we relied on a *’stage-span’* window σ, so that individual cells will be withdrawn from the pooled system σ *t.u.* after completion of TCR*β* recombination. In this context, σ, which is not an additional parameter in the model (whatever setting value, the whole dynamics will be respected), simply prevents cell accumulation. Importantly, in the constant inflow situation, and from the steady state asymptotic regimen onwards, ratios of the various TCR*β* genotype outcomes are mathematically identical to those in the synchronous case described above [*SI Appendix, S1*/equations (15) and (16)].

In summary, considering the synchronous or asynchronous case without bias, the probability distributions ℙ_*i*_(*t*_*i*_ = *t*), defined along with equation (1), determine (*via* the spectrum of *δ* = *t*_2_ −*t*_1_ values) the cellular profiles in terms of TCR*β* genotype at any time *t*. In particular, for given values of [*τ*, *ρ*], final cell profiles (of which there are five -*g*_3_, *g*_5_, *g*_6_, *g*_7_ & *g*_9_-that are divided across the three generic classes *ψ*_1_, *ψ*_2_, and *ψ*_3_, as described above) depend only on these two related sets of scattered values (*SI Appendix, S1*/*Suppl. Table 1*).

### Calibration of the model system reveals approved values of TCR*β* 60/40 ratio

We first proceeded with model calibration, to ascertain whether our model system may be solved such that predictions of AE measurable observables can be obtained. In the situation where *p*_1_=*p*_2_=*p*, equation (13) of the model may be used to delineate confidence intervals for parameter values [*p*, *τ*, *ρ*], based on experimental data. Previous studies of the TCR*β* sequences from wildtype (wt) mouse T-cells or T-cell hybridomas yielded quantitative information on cell populations displaying only three out of the five potential TCR*β* final states, namely [*g*_3_, *g*_5_ and *g*_9_] (Aifantis et al., 1997) or [*g*_3_, *g*_5_ and *g*_7_] (Senoo et al., 2003; Khor and Sleckman, 2005) (*SI Appendix, S1*/*Suppl. Table 2*). In equation (13), *F* is a strictly monotonic function of *p* for any fixed *α*, whereas each of the two *g*_9_ (VDJ+/VDJ+) and *g*_7_ (VDJ+/GL) fractions (which we named *u* and *υ*, respectively) is found in only the *ψ*_1_ or *ψ*_3_ subclass, respectively. Consequently, provided that *ρ* is set first (without loss of generality we set *ρ* = 1 *t.u.*), *p* can be calculated from *u*; then, *τ* can be calculated from *υ*. From data in (Aifantis et al., 1997), a mean estimate for *u* was 1.8%, corresponding to *p*_*wt*_=0.0606. From data in (Khor and Sleckman, 2005; Senoo et al., 2003), estimates for *υ* vary from 1% to 2.6%, thereby setting 64 ≥ *τ*_*wt*_ ≥ 48 (when *p*_*wt*_=0.0606). In using these boundary values for testing simulations, we verified that the resulting [VDJ+/DJ (or GL)] to [VDJ+/VDJ-] cell ratio falls within the range of experimentally-observed distributions [*e.g.* 54.6/45.4 when *p*_*wt*_=0.0606 (*u*=1.8); *τ*_*wt*_=55 (*υ*=1.87); (Mostoslavsky et al., 2004; Brady et al., 2010a)]. Importantly, we further verified that moderate variations in the parameters’ values still yield genotype outputs compatible with TCR*β* AE hallmarks, a pledge of stability of the model system (*SI Appendix, section S3*).

### Model studies predict elements of robustness of the TCR*β* AE process

To further characterize the properties of our model, we simulated the evolution of TCR*β* genotypes within synchronous cells displaying distinct sets of *p* values (Fig. 3), including those in the wt case [row A, (*p*_1_=*p*_2_=*p*_*wt*_=0.0606)] or those in situations where one or both *p* were either decreased or increased relative to the *p*_*wt*_ baseline [rows B, (*p*_1_=*p*_*wt*_, *p*_2_=0.01); C, (*p*_1_=*p*_2_=0.01) and D, (*p*_1_=*p*_*wt*_, *p*_2_=0.4); E, (*p*_1_=*p*_2_=0.4)]. We performed similar simulations of asynchronously developing cells considering the (*p*_1_=*p*_2_=*p*_*wt*_) condition and two types of influx supply dynamics, either constant or periodic (Fig. 4A/B). Thus, we verified the predicted synchronous/asynchronous equivalence for final subpopulations in the case of constant influx (*e.g.* far right panels in Fig. 3A/4A). Likewise, even though stable ratios are not permanently conserved in the periodic influx situation (Fig. 4B; observed oscillations here reflect on and off periods in flow supply), averaging these quantities over a whole oscillation period *in fine* yields distributions similar to those in the case of constant influx (Fig. 4C). In short, insofar as *g*_3_, *g*_5_, *g*_7_, and *g*_9_ populations are considered (in practice those that are commonly accessible to the experimentalist), synchronous and asynchronous regimens using the same *u*-and *υ*-based parameterization of the modeling system lead to identical distributions-and fit those found in *bona fide* biological samples of developed T-cells [*SI Appendix, S1*/*Suppl. Table 2*; (Mostoslavsky et al., 2004; Brady et al., 2010a)]. Only in unusual situations where the inflow/outflow may adopt a prolonged period of continuously increasing or decreasing magnitude [*e.g.* linked to aging (Balomenos et al., 1995)] might the end distributions be modi1ed. However, we note that the synchronous regimen provides a more complete picture of evolutionary dynamics, while the asynchronous regimen may cause information loss in summing up transient cells from different developmental stages.

**Figure 3.**
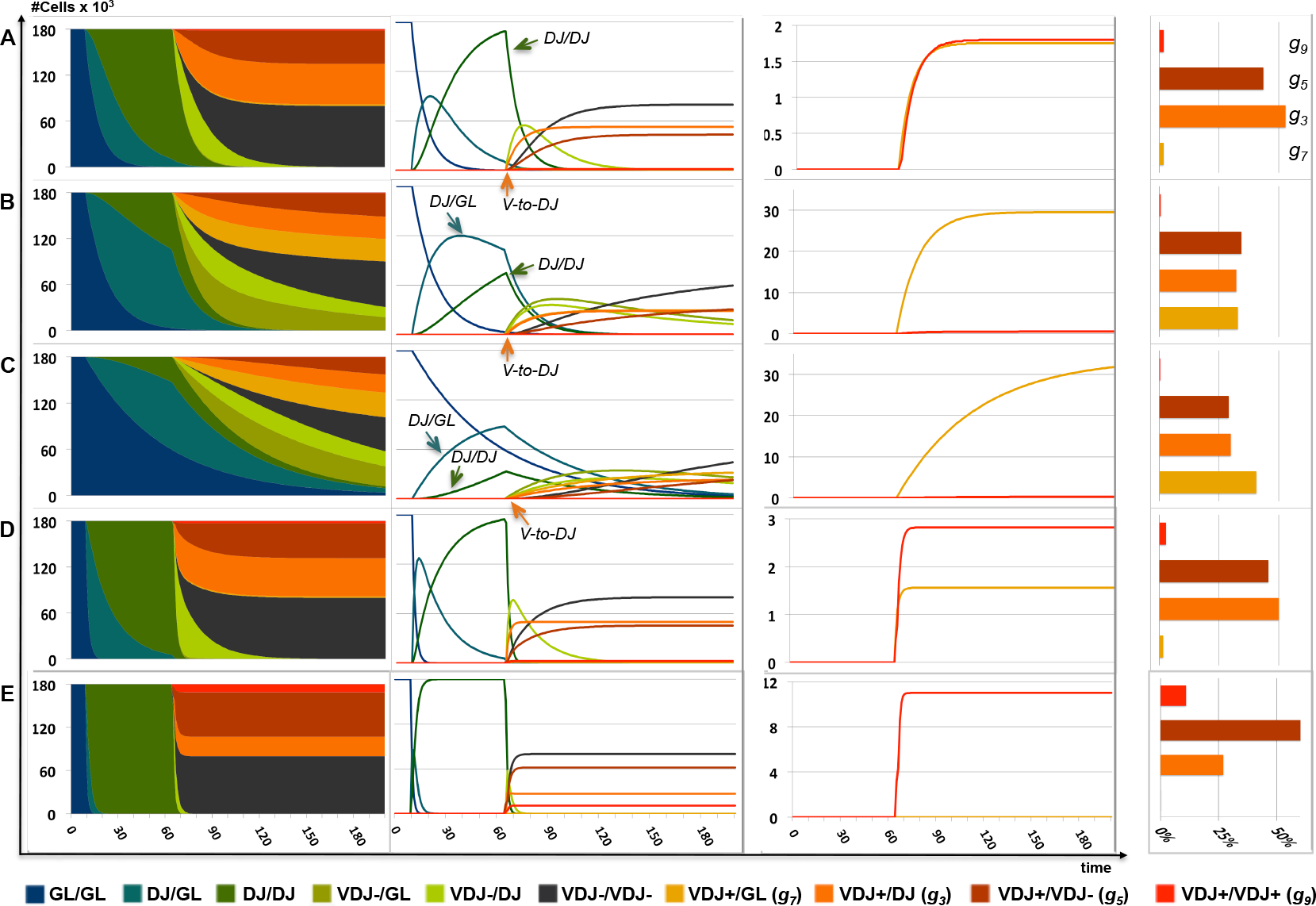
Dynamics of TCR*β* gene recombination considering synchronously developing T-cells. The evolutions in the distribution of the *g*_0_-*g*_9_ TCR*β* allelic genotypes are shown in the cases of cell populations where (**A**) *p*_1_=*p*_2_=*p*_*wt*_; (**B**) *p*_1_=*p*_*wt*_, *p*_2_=*p*-; (**C**) *p*_1_=*p*_2_=*p*-; (**D**) *p*_1_=*p*_*wt*_, *p*_2_=*p*+; and (**E**) *p*_1_=*p*_2_=*p*+, with *p*_*wt*_=0.0606, *p*−=0.01 and *p*+=0.4. The remaining parameters are identical in these five situations (*τ*=55; *t*_*x*_=10; *ρ*=1). The two left panels show distinct representations (in cumulative and separate manners, respectively) of identical dynamics, using a common scale of cell numbers, as indicated on the left. The middle right panel is a magnification of the curves corresponding to the separate VDJ+/VDJ+ and VDJ+/GL genotypes only. The bars on the far right panels indicate the percentage distribution of each final subpopulation *g*_3_ (VDJ+/DJ), *g*_5_ (VDJ-/VDJ+), *g*_7_ (VDJ+/GL) and *g*_9_ (VDJ+/VDJ+) at the end of the depicted dynamics (200 *t.u.*). Arrows on the dynamical curves in (**A-C**) point to the maximum values of the DJ/GL and DJ/DJ genotypes (dark green arrows), and to the initiation of V-to-DJ rearrangement (orange arrows).

To reveal the dynamical attributes of the model, we next examined the simulated trajectories depicted in Fig. 3. In the wt situation, TCR*β* final states compatible with experimental margins are all handled relatively shortly after initiation of the dynamic process, without the need for any additional parameter (Fig. 3A; asymptotic regimens beginning at *t* = 90-120 *t.u.*). In this setting, the DJ/DJ genotype has mostly reached its maximal value (and consequently few alleles remain in the GL configuration) when the first attempt for V-to-DJ rearrangement takes place (arrows); this accounts, with minimal constraint, for the general view that most developing T-cells have achieved D-to-J recombination on the two TCR*β* alleles before the initiation of V*β*-to-DJ*β* rearrangement (Mostoslavsky et al., 2004; Brady et al., 2010a). However, the dynamics and genotype distributions change as *p* decreases relative to the *p*_*wt*_ baseline at one or both allele(s). Then, the distributions-when compared to the wt alleles-display slower kinetics and equivalent or even higher fractions of transient DJ/GL *versus* DJ/DJ genotypes at the time of a first V-to-DJ rearrangement (Fig. 3B-C, left two panels). In both cases, this was followed by increased representation of VDJ+/GL (*g*_7_) cells and, conversely, by decreased fractions of both VDJ+/DJ (*g*_3_) and VDJ+/VDJ-(*g*_5_) cells and near-complete absence of VDJ+/VDJ+ (*g*_9_) cells (right two panels). Overall, this implies that in these two instances, TCR*β* AE would be enforced while the system’s productivity (in terms of kinetic efficiency to yield comparable amounts of productively rearranged cells) would be reduced. Finally, the whole dynamics appear accelerated as *p* increases on one or both alleles, with, in either condition, the somewhat earlier implementation of asymptotic profiles (Fig. 3D-E). However, in contrast to the above findings, eventual effects on cell distribution pertain only to the symmetrical (*p*_1_=*p*_2_) situation, which solely displays increased VDJ+/VDJ+ (*g*_9_) cells, near-complete absence of VDJ+/GL (*g*_7_) cells, and an inversion of the VDJ+/DJ (*g*_3_) to VDJ+/VDJ-(*g*_5_) ratio. This suggests that an additional, unanticipated function of *p* in this system is to act as a point of regulation to ensure robustness of the AE process.

**Figure 4.**
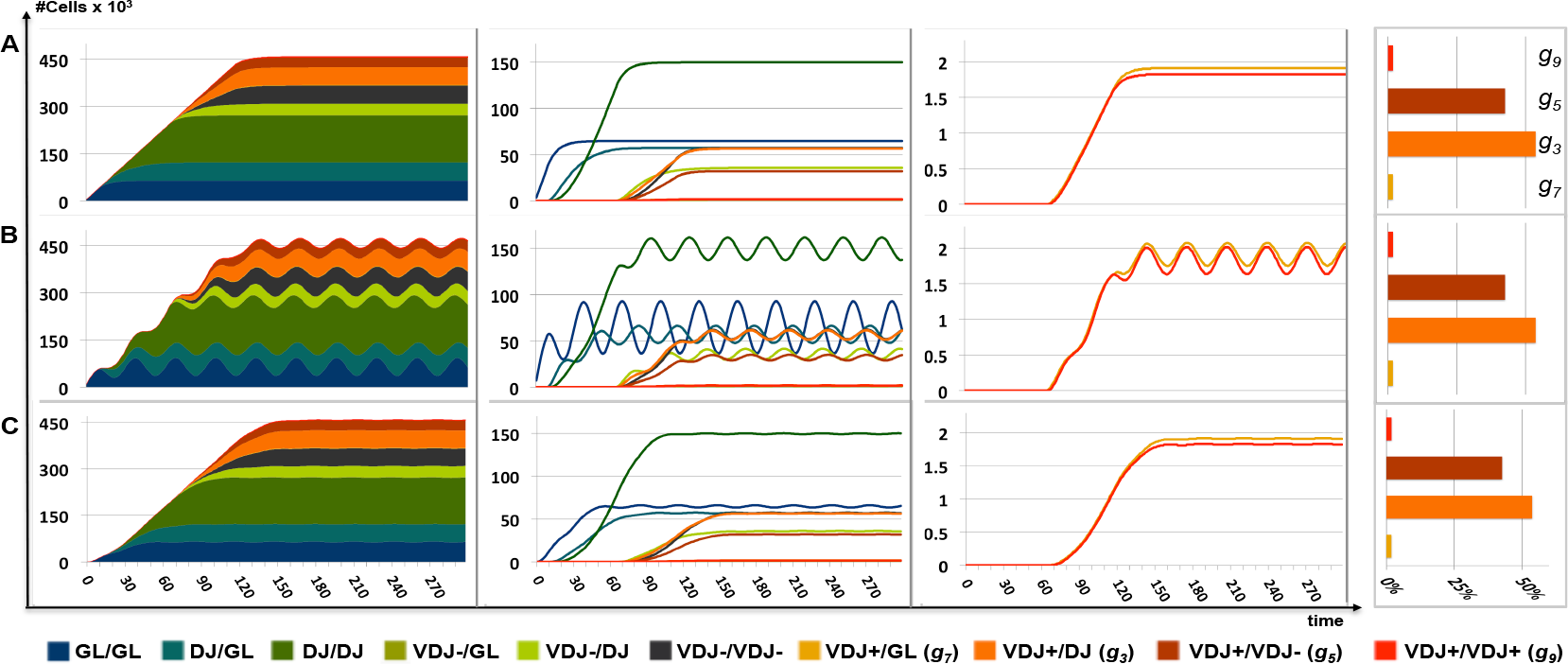
Dynamics of TCR*β* gene recombination considering asynchronously developing T-cells. The evolutions in the distribution of cell populations of the *g*_0_-*g*_9_ TCR*β* genotypes are shown in the situation of constant (**A**) or periodic (**B**) influx supply dynamics; both with *σ* = 50 *t.u.* Parameter values are the same as in Fig. 3A. (**C**) corresponds to the periodic influx situation [as in (**B**)], following integration of distribution mean values over a sliding window of 30 *t.u.* (theoretical oscillation period = 30.4 *t.u.*). The bars on the far right represent the distributions of the indicated cell subsets, as integrated at the end of the simulation period (300 *t.u.*; once their *’stage-span’* window *σ* has expired).

To thoroughly explore the effects of *p* value variation on the yield of allelically included cells, we established the plot *u*(*p*_1_, *p*_2_). Thus, *u* values mostly display a growing series of ‘L-shaped’ isocurves (Fig. 5). Appreciable expansion of *u* [where VDJ+/VDJ+ cells are more than around 4-5% of mature T-cells, up to the theoretical 20% max. when AE is nil] requires that *p* be increased on both alleles (note that, conversely, AE is ultimately enforced when either one or both *p* is/are decreased, as predicted). In the common situation of symmetric *p* incrementation, *u* reads along the diagonal *p*_1_=*p*_2_. In asymmetric cases [*p*1<*p*2 (resp. *p*2<*p*1)], however, *u* displays ‘off-diagonal’ entries that hold within values relatively close to the corresponding lower *p*_*i*_ on the diagonal, and maximum values (and therefore extent of allelic inclusion) that are restrained [Fig. 5, one example (*p*_1_=0.3) is illustrated].

**Figure 5.**
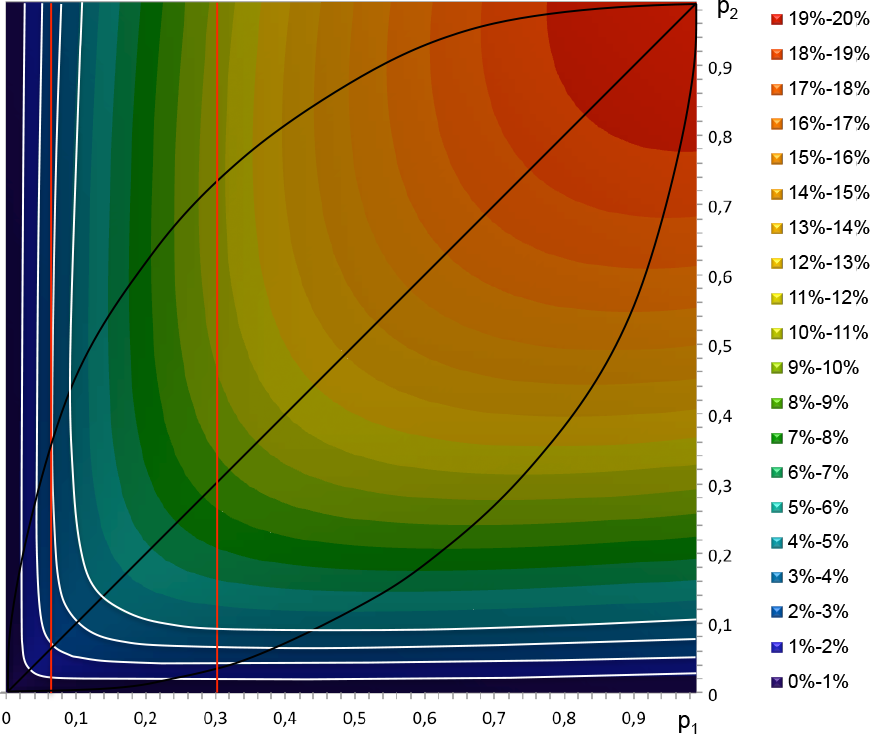
Plot of *u* as a function of parameters *p*_1_ and *p*_2_. White curves delineate constant values of *u* (= % of allelically included VDJ+/VDJ+ cells), from *u*=1% to *u*=4%. Black curves delimitate the values of *p*_2_ that maximize *u* as a function of *p*_1_ (top left curve) and, conversely, those of *p*_1_ that maximize *u* as a function of *p*_2_ (bottom right curve). The diagonal is for *p*_1_=*p*_2_=*p*. The vertical red lines indicates the curves′ intersection with the *p*_1_=*p*_*wt*_=0.0606 value estimated in this study (after optimization-see *SI Appendix, section 3*), as well as with the *p*_1_=0.3 value.

Likewise, when *e.g. p*_1_ (resp. *p*_2_) is kept constant (*p*_1_=*p*_*wt*_), and *p*_2_ (resp. *p*_1_) aug-ments progressively, *u* also shows little variability-initially increasing slightly and then decreasing (Fig. 5, left red line)-so that the fraction of VDJ+/VDJ+ cells is maintained within an acceptable range regarding AE profiles. In the *SI Appendix, section S4*, we propose an explanation for these surprising behaviors of robustness, based on the compensation effect of *δ*s’ values between cells that first initiate rearrangement of either allele 1 or allele 2. We conclude that the simulations we demonstrate here suggest that our model is a versatile system able-for the first time to our knowledge-to satisfy, with minimal constraints, known features of, and long-lasting questions relating to, AE and its initiation phase at the TCR*β* locus (see introduction). Furthermore, the model appears to comprise key elements of stability and robustness, helping to optimally manage the dual objective of promoting independent dual allele usage and uneven development of allelically excluded/included cells. This may be essential, considering a system in which mono-allelic expression primarily depends on imprecise DNA recombination events that frequently results in non-functional CJs.

### Model studies predict measurable outcomes from DJ and VDJ knock-in studies

Two studies that analyzed strains of knock-in (KI) mice carrying either a rearranged DJ or a rearranged VDJ gene segment on one or both TCR*β* alleles (Carpenter et al., 2009; Brady et al., 2010b) were of particular interest for further testing our model, in which asymptotic equilibria depend on relative timespans, including those required to achieve D-to-J (*t*_*x*_) and V-to-DJ (*t*_*y*_ = *t*_*x*_ + *τ*) rearrangements. Both studies established end distributions of the VDJ/DJ and VDJ/VDJ configurations of TCR*β* alleles using T-cell hybridomas from KI animals. We relied on these data, in particular those derived from heterozygous T-cells in which wt and mutant alleles can compete simultaneously, to challenge the fit of our model after either (*t*_*x*_) or (*t*_*x*_ + *τ*) were set to 0 for one allele, binding the other allele to wt values. All other parameters were maintained as defined above. In addition, the initial cell population was set to a DJ/GL or VDJ+/GL configuration (rather than GL/GL, as in the wt situation).

Based on the analysis of 247 hybridomas harboring one DJ*β*-preassembled complex, Carpenter *et al.* reported a distribution of 58% VDJ/DJ and 42% VDJ/VDJ, providing evidence of a two-step recombination process that is dispensable for enforcement of TCR*β* AE (Carpenter et al., 2009). Remarkably, simulations using our model system yielded comparable values of 57% VDJ/DJ and 43% VDJ/VDJ (Fig. 6A; *SI Appendix, S1*/*Suppl. Table 3*), showing that our modeling design of AE is consistent with these biological figures. Moreover, in VDJ+/[DJ or GL]-predicted cells, with a ratio of VDJ joints at the (DJ)-and (GL)-initially configured allele of 74% to 26%, respectively, the simulations reproduce the observation that the majority of the corresponding hybridomas contain V*β*DJ*β* rearrangements on the KI allele [69% *versus* 31%, as reported; Carpenter et al. (2009)]. In fact, our dynamical view of allele asynchrony from locus-activating events onwards offers an explanation for this phenotypic bias: the mutant allele, when ranking second in starting the recombination process with a disadvantaged *δ* relative to its wt homologue, nevertheless has the opportunity to achieve a VDJ+ joint first for each *δ* < *t*_*x*_, thus adding to the number of cases where it starts and finishes first. This competitive advantage of the mutant allele is reflected in the reduced time needed for VDJ/[DJ or GL] configurations to emerge in the simulations, and by the biphasic dynamics of the VDJ+/GL genotype (Fig. 6A; to be compared with the corresponding data in Fig. 3A).

An analysis of 171 T-cell hybridomas harboring one functionally preassembled V*β*DJ*β* complex showed a distribution of 5.3% V*β*DJ*β*-rearranged and 26.9% DJ*β*-rearranged wt alleles, respectively (Brady et al., 2010b). This distribution, showing a significantly reduced DJ fraction, is surprising, as former studies have found that ectopic expression of a VDJ+ exogenous transgene, though inhibiting the endogenous V-to-DJ rearrangement, had no effect on the course of D-to-J recombination [Mostoslavsky et al. (2004); Cedar and Bergman (2008); Brady et al. (2010a); Vettermann and Schlissel (2010); and thus it has generally been accepted that TCR*β* AE impinges exclusively on V gene recombination]. However, these conflicting results can be better understood using a dynamical model system like ours, which encompasses all stages of the recombination process (with the reasonable assumption that ectopic *versus* site-specific genomic integrated VDJ+ inserts behave differently in terms of timing in developmental expression). Indeed, resulting simulations from our revised model (*t*_*x*_= / *τ*=0 at one allele, which is also considered to be functional in all cells, and not just one third of them, throughout the simulation period) retrieved values of 0.94% (V*β*DJ*β*) and 28.43% (DJ*β*) on the wt alleles (Fig. 6B; *SI Appendix, S1*/*Suppl. Table 3*). It is noteworthy that the relatively low numbers of experimental samples (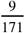 and 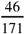 hybridomas, respectively) may not be without statistical impact on the confidence intervals for the biological data, especially with respect to the V*β*DJ*β* fraction. This may partly account for the variation between values reported in (Brady et al., 2010b) and our simulation results. Furthermore, the KI mice used in this study harbor TCR*β* alleles with duplicated D*β*1-J*β*1 and D*β*2-J*β*2 clusters on the wt allele and a D*β*2-J*β*2 cluster downstream of the inserted segment on the mutant allele. As parallel analyses implied that DJ*β*2 complexes are used in further V-to-DJ rearrangement on the mutant allele in ~3% of homozygous hybridomas (Brady et al., 2010b), one may indeed expect a slightly increased fraction of VDJ+ wt alleles in this case of D-J cluster duplication, *i.e.* following cell rescue as also predicted from our current modeling views (*SI Appendix, section S5*). Overall, we therefore conclude that our model system is not only able to satisfactorily simulate published data, but also able to provide a straight forward explanation for the observations that KI of a DJ*β*-rearranged segment does not impact upon AE at the T-cell population level, yet results in an increased frequency of VDJ+ rearrangements at this site (Carpenter et al., 2009). Furthermore, the model system also corroborates data showing that KI of a VDJ+ preassembled segment does impact, to varying degrees, on both V-to-DJ *and* D-to-J rearrangement outcomes at the opposite wt allele (Brady et al., 2010b). Considering the overall picture drawn by our simulations, the above statements also suggest that the stochastic events that open the TCR*β* alleles for recombination and/or expression are independent of the rearrangement status (GL, as in the wt allele; DJ*β* or V*β*DJ*β*, as in the KI allele). We note that one hallmark common to all these alleles, regardless of their configuration, consists of the presence of the enhancer E*β*, in association with either the promoters pD*β* or V*β* and ranking architectural, long-range tethering elements [*e.g.* Majumder et al. (2015)]. Recent evidence suggests that critical shifts during lymphoid cell development are accompanied by large-scale changes in epigenetic markers associated with enhancers, promoters and architectural elements of cell-lineage-specific genes, including antigen receptor genes (Benner et al., 2015). In line with this and our modeling precepts, we propose that chromatin opening of these various alleles proceeds coincidentally (though not necessarily simultaneously)-mostly involving E*β*-driven probabilistic outputs and functional interactions between these various *cis*-regulatory elements in the 3D nuclear context-to begin the course of TCR*β* gene expression, recombination and appended AE profiles, as observed in the corresponding T-cells.

**Figure 6.**
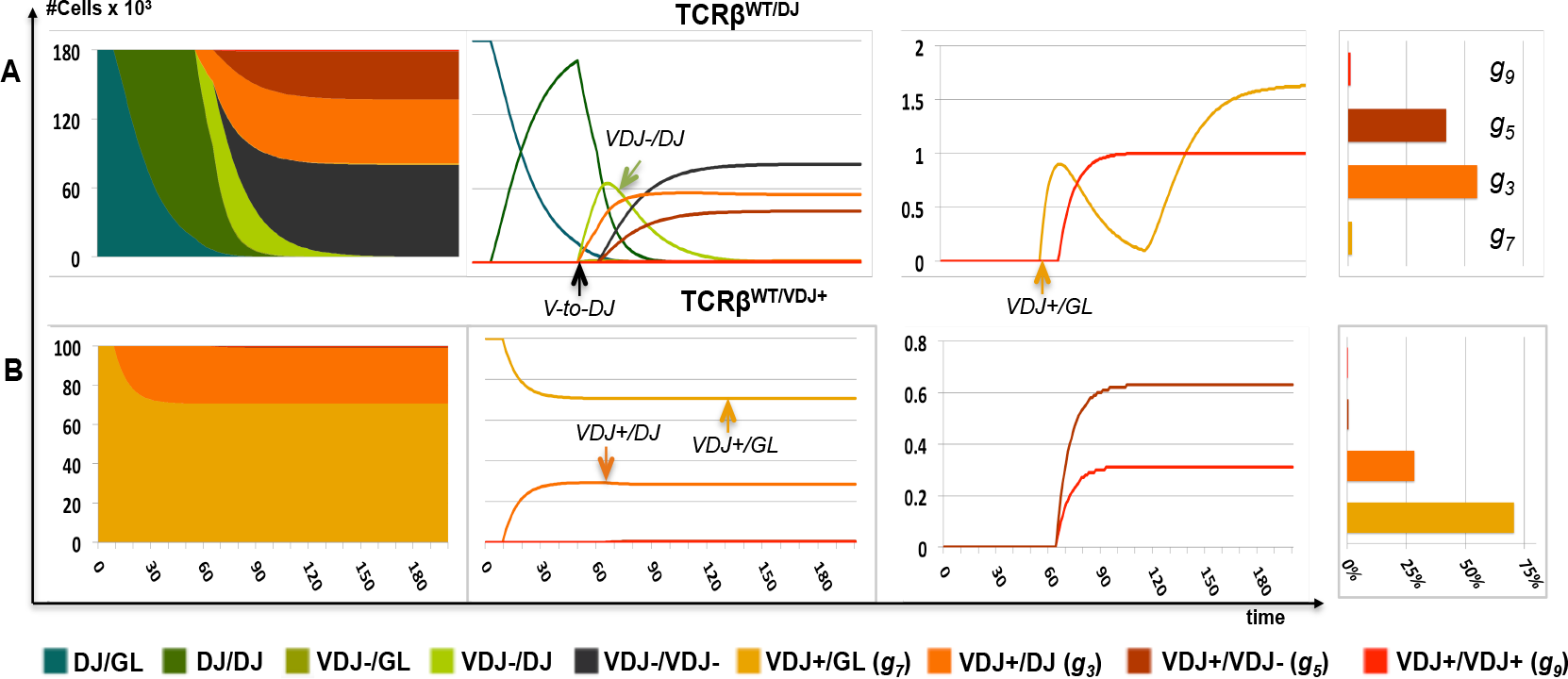
Box initially at rest on sled sliding across ice.

Reproducing data from knock-in (KI) DJ-rearranged and VDJ-rearranged TCR*β* alleles. Evolution in the distribution of the *g*_1_-*g*_9_ TCR*β* allelic genotypes considering developing T-cell populations carrying either one wt allele and one KI DJ*β*-rearranged allele [TCR*β*^*wt*/*DJ*^(**A**)] or one wt allele and one KI functionally-rearranged V*β*DJ*β* allele [TCR*β*^*wt*/*V DJ*+^(**B**)]. Legend is as in Fig. 3, except that the middle right panel in (**B**) shows a magnification of the curves corresponding to the separate VDJ+/VDJ-(*g*_5_=0.63%) and VDJ+/VDJ+ (*g*_9_=0.31%) genotypes (total V*β*DJ*β*=0.94%). (**A**): *p*_1_=*p*_2_=0.0606; *t*_*x*_=0 at one allele; (**B**): *p*_1_=*p*_2_=0.0606; *t*_*x*_=0 and *τ*=0 at one allele. All others parameters as defined in the Results. According to model implementation, the TCRTCR*β*^*wt*/*DJ*^ and TCRTCR*β*^*wt*/*V DJ*+^distributions lack the GL/GL (*g*_0_) configuration. The biphasic behavior of the VDJ+/GL genotype in (**A**) reflects the early appearance of VDJ+/GL cells because of the dynamical advance of the KI DJ allele (V-DJ rearrangement on this allele), followed by their progressive evolution into VDJ+/DJ cells (D-J rearrangement on the wt allele) and, eventually, by their regular emergence within the kinetic system.

### Model studies predict observable TCR*β* repair foci and system-relevant timings

We next wished to examine whether a better temporal picture of our model could be obtained from experimental results. In the absence of real-time insights into sequential rearrangements at dual TCR*β* alleles in single DN cells, we considered a recent study that used 3D immuno-fluorescence *in situ* hybridization (FISH) to visualize newly generated RAG-dependent DSBs at TCR loci based on the accumulation of the DNA-repair protein 53BP1 (Chan et al., 2013). However, this study reported a low ratio of 53BP1+ TCR*β* foci in DN thymocytes (<10%), which, in light of the recurrent positioning of TCR*β* alleles at the nuclear periphery (Schlimgen et al., 2008), has been taken to suggest that such alleles are mostly inefficient in their ability to undergo V(D)J recombination. Yet, when regarded from a dynamic angle, immuno-FISH data and its significance in terms of recombination frequency must be weighted relative to mean-time values of both cell residency within the system and length of focus detection (for more details, see *SI Appendix, section S6*). On this basis, new calculations again proved the predictive power of our model system in this specific instance and, furthermore, allowed us to predict real-time features of the system. Initially, considering *(i)* reasonable adjustments regarding lifespan kinetics of developing DN cells (Vasseur et al., 2001) as well as of 53BP1+ foci at sites of DNA DSBs (Asaithamby and Chen, 2009), and *(ii)* anticipated scenarios of cell developmental histories and their distributions into distinct *ψ*_1_, *ψ*_2_, and *ψ*_3_ subsets, as a first approximation we obtained likelihoods of 9.27% and 0.53% for single and dual observable foci, respectively (*SI Appendix, S6A*), which closely correlates with the corresponding 9.4% and 0.4% reported in (Chan et al., 2013). Building on these assets, we then took advantage of the latter experimental values to determine-following mathematical refinements aimed at more precisely tracking RAG-dependent DSB kinetics throughout the *ψ*_1_, *ψ*_2_, and *ψ*_3_ subsets-realistic estimates for mean residence time and time unit in the model system [RT = 89*h*; *t.u.* =1.18*h*; *SI Appendix, S6B*, equations (17)-(20) and Fig. S6-2]. This offered the opportunity for a direct assessment of typical time-frames in the system (Table 1; *SI Appendix, S6B*, Additional comments #1 and #2). Importantly, we note that the former two estimates of RT = 89*h* and *t.u.* =1.18*h* are consistent with experimentally derived approximations of the duration of the DN2-DN3 developmental window (Vasseur et al., 2001) and of total timings supposedly integrated by *ρ* = 1 *t.u.* (*SI Appendix, section S7*), respectively. We further note that temporal interconnections point to the effectiveness of the system’s operation. Notably, median values of initiation time [mIT; 9 *t.u.*/10.6*h*] are sufficiently lower, and interallelic time gap [m*δ*; 11 *t.u.*/13*h*] are sufficiently higher, compared to mean residence time [RT; 75 *t.u.*/89*h*] and feedback inhibition [*ρ*; 1 *t.u.*/1.18*h*], respectively, to bolster both bi-allelic usage and hence cell productivity, and the preponderance of allelically excluded individuals. Finally, it is now obvious that these values better fit the above immuno-FISH data than those derived from previous Markov-chain-based simulations, in which stochasticity mostly pertains to the V*β*-to-DJ*β* rearrangement step [(Farcot et al., 2010); *SI Appendix, S6B*, Additional comments #3 and #4]. When also considering that, although not further discussed in their article, data reported in (Chan et al., 2013)-in particular the low ratio of dual/total 53BP1+ foci and corollary of sparse contemporary DSBs-imply that both steps of TCR*β* recombination are likely to occur asynchronously, we contend that these additional simulation results provide further compelling evidence supporting the consistency of our model.

**Table 1.** Relative and estimated absolute time values for mean residence time inside the system [RT]; median value of initiation time [mIT] (which we define as the time when 50% of the cells have initiated a first D-to-J rearrangement, and which therefore re2ects the effixsciency of the recombination initiation phase); median value of interallelic time gap *δ* [m*δ*]; and dwell-time values of D-to-J rearrangement [*t*_*x*_], V-to-DJ rearrangement [= parameter *τ*], and feedback inhibition [= parameter *ρ*].

| | [RT] | [mIT] | [mδ] | $t_x$ | $\tau$ | $\rho$ |
| --- | --- | --- | --- | --- | --- | --- |
| Relative Time ($t.u.$) | 75 | 9 | 11 | 10 | 55 | 1 |
| Absolute time (hours) | 89h | 10.6h | 13h | 11.8h | 65h | 1.18h |

## Discussion

We created a mathematical model in order to test the hypothesis that cell-to-cell interallelic variability during the initiation phase of TCR*β* gene recombination can account for typical AE profiles at this locus. The model is based on a limited set of parameters-*p*, *τ*, and *ρ*, which define critical time intervals during the recombination process. This work validates our hypothesis in the context of both synchronous and asynchronous flow-dependent behaviors of developing cells. Simulations under diverse allelic conditions, to mimic experimental studies, yielded data that underscore the predictive power of the model.

At V-D-J structured loci, the initiation phase of AE (allele asynchrony) is usually viewed in the context of V-to-DJ assembly, because of observations that dually DJ rearranged *Igh* or TCR*β* alleles are present in nearly all pre-BCR/pre-TCR selected cells. Since those findings, however, signs of an unforeseen dissociation of D-to-J recombination on opposite alleles have surfaced (Born et al., 1988; Akamatsu et al., 2003; Dudley et al., 2003; Bonnet et al., 2009). Our simulations reconcile all these data, at lower regulatory cost, as they support a dynamical cell-system in which allele asynchronicity generated by probabilistic events prior to D*β*-to-J*β* rearrangement can account for the observed distribution of TCR*β* genotypes, while also highlighting that the DJ/DJ prevalence can peak prior to the first V-to-DJ joining attempt. Moreover, the model provides rational explanations of the antipodal observations of V*β* gene AE, DJ*β* joining ubiquity, and seeming scarcity of RAG-mediated DSB foci at TCR*β* alleles (Chan et al., 2013). In fact, the model provides these explanations without suggesting that the TCR*β* alleles are inefficient in their ability to undergo V(D)J recombination (after all, nearly all cells experience dual allele activation/rearrangement at some point in their own development), eventually reconciling stochasticity in recombination with developmental cell productivity [see *e.g.* Gorman and Alt(1998)]. Finally, the model system also presents a broader interpretation of various genetic mutations known to affect AE in T lymphocytes (*SI Appendix, S7B*).

Under this model, asymptotic distributions of TCR*β* genomic profiles in T-cells split into three subgroups (*ψ*_1_, *ψ*_2_, and *ψ*_3_), depending on the relative values between the interallelic time gap *δ* and parameters *τ* and *ρ*, with *δ* > *ρ* as a key precept to enforce AE. From a biological perspective, we believe that *δ* values in DN cells primarily rely on the asymmetric FPT timing of combinatorial, activating epigenetic changes. These [changes in histone marks from *e.g.* H3K9me2 to H3K4me3, Pol II-generated short-RNA transcripts, CpG demethylation, mobilization of nucleosomes; Morshead et al.(2003); Sikes and Oltz (2012); Kondilis-Mangum et al. (2010); Zacarias-Cabeza et al. (2015)] would be triggered on opposite D*β*-J*β* loci by the concerted activity of adjacent *cis*-regulatory elements, whose combinatorial stochastic outputs would thereby be integrated by geometric distributions dependent on parameters *p*_*i*_. Schematically, primary changes may possibly initiate at E*β* through the recruitment of a discrete set of master transcription factors and/or co-activators/epigenetic modulators, and subsequently be enforced and regionally spread *via* E*β*-pD*β* driven and inter-nucleosome cooperative interactions (Sikes and Oltz, 2012; Zhang et al., 2014). This in turn would enable the segregation from the suppressive nuclear periphery, the structuring of the TCR*β* recombination center, and prime D-to-J rearrangement. Longer term epigenetic and structural reorganization, initially insulated within the D-J-C domains, would also involve-in our view, as a ‘canalized’ process (see below)-the V*β*-containing domains propelling *in 1ne* their folding into the DN-specific 3D-architecture wherein topological confinement permits V-to-DJ rearrangement (Majumder et al., 2015; Hu et al., 2015). Accordingly, the *δ* interallelic hallmarks in individuals cells would grossly be maintained up to this phase. In practice this would be up to the point of ‘prime-encounter’ and ensuing recombination between V and DJ elements (Lucas et al., 2014), on first-and second-rearranging alleles, respectively (unless the process is interrupted by *ρ* timed feedback). Without focused experimental clues on dynamical (epi)genome evolution, at separate TCR*β* alleles, in statistically significant numbers of single DN developing cells, we acknowledge that this overall scenario remains hypothetical. Still, because our ‘few-parameter’ modeling study demonstrates sufficient robustness under various allelic conditions, it is worth considering this possibility, given the remaining mysteries that surround the initiation phase of AE.

Recombination of genomically distant V and DJ segments requires their juxtaposition within merged CTCF-binding element (CBE)-flanked loop domains (Majumder et al., 2015; Hu et al., 2015). Underlying principles that direct these events can be accommodated by our dynamical model if considered as part of a hierarchical progression, in which allele-and cell-specific kinetic fluctuations generated during this step (united under parameter *τ*) remain limited with respect to those responsible for prime asynchronicity (note that we thus do not consider that all these later events are homogeneously invariant at all TCR*β* alleles). It is now recognized that early developing cells could have a relatively high stochastic variation across the genome compared to more differentiated cells, with epigenetically regulated variability impinging on early developmental plasticity, followed by less variable, canalization periods of development to produce consistent phenotypic outcome(s) (Pujadas and Feinberg, 2012). In these contexts, earlier views of autonomous blocks in V-to-DJ recombination, largely from studies involving the *Igh* locus (Corcoran et al., 1998)-when examined with the benefit of hindsight-appear to better support a scenario whereby the activation of the two regions is not necessarily uncoupled (Fuxa et al., 2004; Malin et al., 2010; Subrahmanyam et al., 2012). Overall, we therefore argue that bound *τ* parameterization (and *ρ*, see below) does not invalidate the meaning of the model, as their dwell-time ± standard deviation values plausibly oscillate over shorter time-ranges when compared to those which lead to allelic differential activation.

Parameter *ρ*, which covers the entire period of time from the expression of a VDJ+ assembled gene to feedback *trans*-inhibition, presumably includes basic cellular processes (transcription and RNA processing, nuclear export and translation of mRNA, and protein folding and delivery), as well as DN cell-specific events (including pre-TCR assembly and ensuing signaling and inhibitory outcomes). Although the precise timelines of these processes are unknown, those steps that have been studied individually (in the case of TCR, Ig, or other gene/protein paradigms) were found to be on the scale of minutes to ≤ 1hr duration (*SI Appendix, S7A*, for further discussion of timings supposedly integrated by *ρ*). This suggests that when applied to the 3-4.5 day period necessary for DN2-DN3 development (Vasseur et al., 2001), these steps normally also have limited impact on the course of the dynamical system process, so that most cells would comply with the *δ* > *ρ* precept.

It is tempting to consider that, providing recalibration, our model system could likewise be tailored to the analysis of recombination dynamics at other TCR or Ig genes, and to the analysis of developmental cell fates in distinct lymphoid lineages. This could offer valuable information on mechanistic similarities/disparities among adaptive immune loci subject or otherwise to AE [Mostoslavsky et al. (2004); Cedar and Bergman (2008); Brady et al. (2010a); Vettermann and Schlissel (2010); NB: depending on the locus (and distinct from the TCR*β* locus), recalibration would firstly involve the adjustment of the probabilistic variable *є* ( from the one-third/two-thirds outcome), Vettermann and Schlissel (2010); and would possibly also require model/parameters′ amendment to integrate the cumulative effects of independently controlled chromatin subdomains within larger loci, *e.g.* distal V domains at the *Igh* locus, Montefiori et al. (2016)]. Beyond these applications, our modeling hypotheses might also be adapted to the study of discrete cell types that employ random monoallelic gene expression at the transcriptional level, independently of the genetic sequence and parental origin (*i.e.* distinct from classical examples of X-chromosome inactivation and genomic imprinting). A growing body of evidence demonstrates that a significant fraction of mammalian autosomal genes are subject to monoallelic expression in different cell types, in an inheritable manner (Eckersley-Maslin and Spector, 2014). Indeed, for some such genes (olfactory receptor genes), a kinetic model assuming a tradeoff between slow activation by step-wise epigenetic events and fast feedback signaling satisfactorily accounts for allelic singularity in expression (Monahan and Lomvardas, 2015).

In summary, we provide an original view of TCR*β* AE as a model system of dynamic distribution, primarily based on noise in the initiation phase of V(D)J recombination. Sufficiently reflective of allelically excluded cells, this unique model is remarkably robust and satisfactorily constrains the generation of allelically included stowaways, with no need for any further sophisticated controls. This property may be of significant value-theoretical frameworks have established the extreme cost of multiplying the sources of signaling events when endeavoring to achieve increased biological accuracy (Lestas et al., 2010).

## Materials and Methods

Detailed modeling procedures, including mathematical formulations [Equations (1)-(16)], model programming and computational properties, as well as the collection of experimental data used for the calibration of the model and in modeling simulation studies, and associated references, are available in the *SI Appendix, section S1*/Methods, *section S6*/[Equations (17)-(20)], and *section S8*/References, respectively.

## Acknowledgments

Work in the PF laboratory is supported by institutional grants from ‘Institut National de la Santé et de la Recherche Médicale’ (Inserm) and ‘Centre National de la Recherche Scientifique’ (CNRS), and by dedicated grants from the ‘Agence Nationale de la Recherche’ (ANR), the ‘Institut National du Cancer’ (INCa), the ‘ITMO Cancer Alliance Nationale pour les Sciences de la Vie et de la Santé’ (AVIESAN) and the ‘Fondation Princesse Grace de la Principauté de Monaco’.

